# End-to-end optimization of prosthetic vision

**DOI:** 10.1101/2020.12.19.423601

**Authors:** Jaap de Ruyter van Steveninck, Umut Güçlü, Richard van Wezel, Marcel van Gerven

**Affiliations:** Department of Artificial Intelligence, Donders Institute for Brain, Cognition and Behaviour, Radboud University, Nijmegen, The Netherlands; Department of Biophysics, Donders Institute for Brain, Cognition and Behaviour, Radboud University, Nijmegen, The Netherlands; Biomedical Signal and Systems, MIRA Institute for Biomedical Technology and Technical Medicine, University of Twente, Enschede, The Netherlands

## Abstract

Neural prosthetics may provide a promising solution to restore visual perception in some forms of blindness. The restored prosthetic percept is rudimentary compared to normal vision and can be optimized with a variety of image preprocessing techniques to maximize relevant information transfer. Extracting the most useful features from a visual scene is a non-trivial task and optimal preprocessing choices strongly depend on the context. Despite rapid advancements in deep learning, research currently faces a difficult challenge in finding a general and automated preprocessing strategy that can be tailored to specific tasks or user requirements. In this paper we present a novel deep learning approach that explicitly addresses this issue by optimizing the entire process of phosphene generation in an end-to-end fashion. The proposed model is based on a deep auto-encoder architecture and includes a highly adjustable simulation module of prosthetic vision. In computational validation experiments we show that such an approach is able to automatically find a task-specific stimulation protocol. The presented approach is highly modular and could be extended to dynamically optimize prosthetic vision for everyday tasks and requirements of the end-user.

## Introduction

Globally, over 30 million people suffer from blindness [1]. For some forms of blindness, visual prosthetics may provide a promising solution that can restore a rudimentary form of vision [2–9]. These neural interfaces can functionally replace the eye with a camera that is connected to the visual system. Although the types of visual prostheses may vary considerably (e.g. in terms of their entry point in the visual system), they share the same basic mechanism of action: through electrical stimulation of neurons in the visual system, they evoke a percept of spatially localized flashes of light, called phosphenes [10, 11]. In particular, visual implants that reside in the primary visual cortex (V1) are reported to have an enormous potential in future treatment of visual impairment [5, 9, 12, 13]. Due to the relatively large surface area, this implantation site allows for stimulation with many electrodes. By selective stimulation and by making use of the roughly retinotopical organization of V1, it is possible to generate a controlled arrangement of phosphenes, in such a way that they may provide a meaningful representation of the visual environment (figure 1) [9].

**Figure 1.**
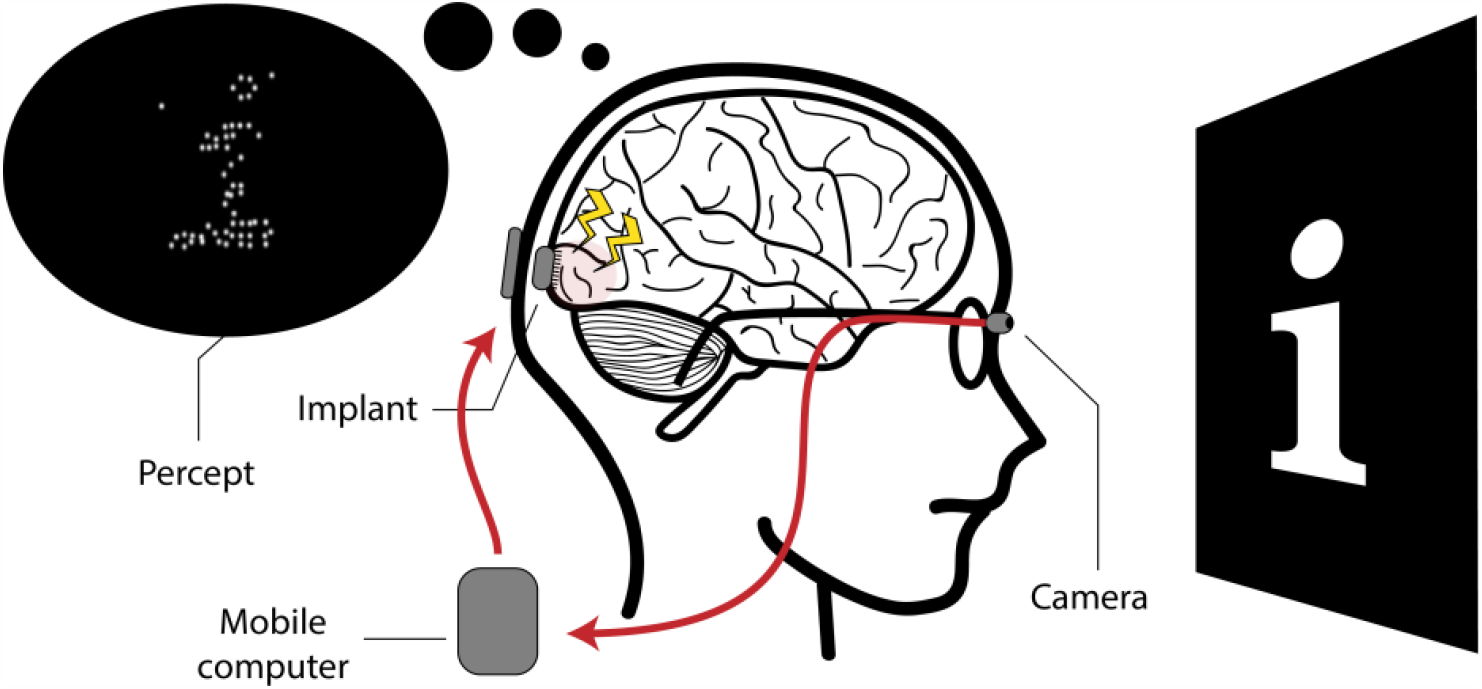
Schematic illustration of a cortical visual neuro-prosthesis. The visual environment is captured by a camera and sent to a mobile computer. Electrodes in the brain implant are selectively activated to stimulate neurons in the primary visual cortex (V1). Making use of the retinotopical organization of V1, a controlled arrangement of phosphenes can be generated to create a meaningful representation of the visual environment.

Compared to normal vision, the percept that can be restored with visual prostheses is very rudimentary and the resolution remains relatively limited, even with relatively high numbers of electrodes (thousands of electrodes) compared to normal vision. The limited amount of information that can be conveyed allows for only selective visualization of the surroundings. Therefore, an important role in the optimization of prosthetic vision will be fulfilled by image preprocessing techniques. By selective filtering of the visual environment, image preprocessing may help to maximize the usefulness and interpretability of phosphene representations. The choice of filtering is non-trivial and the definition of useful information will strongly depend on the context. Therefore, the implementation and optimization of image preprocessing techniques for prosthetic vision remain active topics of scientific investigation.

In previous work, various preprocessing techniques have been tested for a variety of applications using simulated prosthetic vision (SPV) [14–17]. Such preprocessing techniques range from basic edge-filtering techniques for increasing wayfinding performance [18] to more sophisticated algorithms, such as segmentation models for object recognition, or facial landmark detection algorithms for emotion recognition [19, 20]. The latter two examples underline the potential benefits of embracing recent breakthroughs in computer vision and deep learning (DL) models for optimization of prosthetic vision.

Despite the rapid advancements in these fields, research currently faces a difficult challenge in finding a general preprocessing strategy that can be automatically tailored to specific tasks and requirements of the user. We illustrate this with two issues: First, prosthetic engineers need to speculate or make assumptions about what visual features are crucial for the task and the ways in which these features can be transformed into a suitable stimulation protocol. Second, as a result of practical, medical or biophysical limitations of the neural interface, one might want to tailor the stimulation parameters to additional constraints. In this paper we present a novel DL approach that explicitly addresses these issues in image preprocessing for prosthetic vision. We use a deep neural network (DNN) auto-encoder architecture, that includes a highly adjustable simulation module of prosthetic vision. Instead of optimizing image preprocessing as an isolated operation, our DL model is designed to optimize the entire process of phosphene generation in an end-to-end fashion [21]. As a proof of principle, we demonstrate with computational simulations that by considering the entire pipeline as an end-to-end optimization problem, we can automatically find a stimulation protocol that optimally preserves information encoded in the phosphene representation. Furthermore, we show that such an approach enables tailored optimization to specific additional constraints such as sparse electrode activation.

## Methods

In this section, we provide an overview of the components of the proposed end-to-end DL architecture. Next, we describe three simulation experiments that were conducted to explore the performance of our model with personalized phosphene mappings, various sparsity constraints and naturalistic visual contexts.

### Model description

The end-to-end model consists of three main components: an encoder, a phosphene simulator and a decoder (figure 2). Given an input image, the encoder is designed to find a suitable stimulation protocol, which yields an output map. The value of each element in this stimulation protocol represents the stimulation intensity of one electrode in the stimulation grid. The encoder follows a fully-convolutional DNN architecture. In all layers, apart from the output layer, leaky rectified linear units are used as the activation function and batch normalization is used to stabilize training (table 1). The Heaviside step function is used as activation function in the output layer to obtain quantized (binary) stimulation values. A straight-through estimator [22] was implemented to maintain the gradient flow during backpropagation.

**Table 1.** Architecture of the encoder component. Conv: convolutional layer; Res: residual block; Pool: max-pooling layer; BN: batch normalization; LReLU: leaky rectified linear unit.

|  | Type | In | Out | Size | Stride | Pad | Normalization | Activation |
| --- | --- | --- | --- | --- | --- | --- | --- | --- |
| 1 | Conv | 1 | 8 | 3 | 1 | 1 | BN | LReLU |
| 2 | Conv+Pool | 8 | 16 | 3/2 | 1 | 1 | BN | LReLU |
| 3 | Conv+Pool | 16 | 32 | 3/2 | 1 | 1 | BN | LReLU |
| 4 | Res | 32/32 | 32/32 | 3/3 | 1/1 | 1/1 | BN/BN | LReLU/LReLU |
| 5 | Res | 32/32 | 32/32 | 3/3 | 1/1 | 1/1 | BN/BN | LReLU/LReLU |
| 6 | Res | 32/32 | 32/32 | 3/3 | 1/1 | 1/1 | BN/BN | LReLU/LReLU |
| 7 | Res | 32/32 | 32/32 | 3/3 | 1/1 | 1/1 | BN/BN | LReLU/LReLU |
| 8 | Conv | 32 | 16 | 3 | 1 | 1 | BN | LReLU |
| 9 | Conv | 16 | 1 | 3 | 1 | 1 | - | Step |

**Figure 2.**
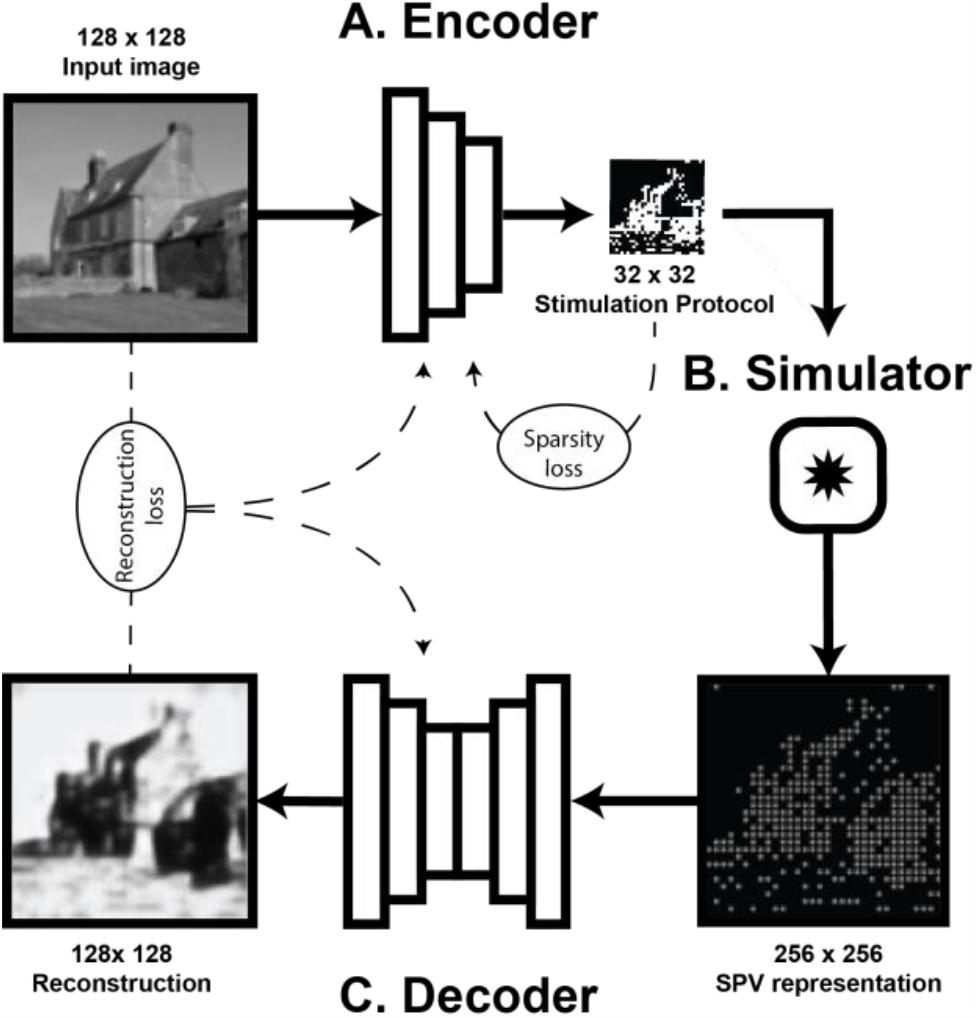
Schematic representation of the end-to-end model and its three components: A. The phosphene encoder finds a stimulation protocol, given an input image. B. The personalized phosphene simulator maps the stimulation vector into a simulated phosphene vision (SPV) representation. C. The phosphene decoder receives a SPV-image as input and generates a reconstruction of the original image. During training, the reconstruction dissimilarity loss between the reconstructed and original image is backpropagated to the encoder and decoder models. Additional loss components, such as sparsity loss on the stimulation protocol, can be implemented to train the network for specific constraints.

In the second component of our model, a phosphene simulator is used to convert the stimulation protocol that is created by the encoder to a SPV representation. This component has no trainable parameters and uses predefined specifications to realistically mimic the perceptual effects of electrical stimulation in visual prosthetics. Phosphene simulation occurs in two steps: first, using a predefined, adjustable, sampling mask, each element in the 32 × 32 stimulation protocol is mapped onto specified pixels of a 256 × 256 image, yielding the simulated visual field. In the current experiments, phosphenes are mapped onto a regular equidistantly spaced grid. Secondly, the obtained image is convolved with a gaussian kernel to simulate the characteristic perceptual effects of electrical point stimulation.

The third component of our model is the decoder. The decoder is an image-to-image conversion model that is based on a residual network architecture [23]. Batch normalization and activation with leaky rectified linear units is implemented in all layers of the model, except for the output layer, which uses sigmoid activation and no batch normalization (table 2). The decoder component is designed to ‘interpret’ the SPV representation by converting it into a reconstruction of the original input. Our end-to-end architecture implements an auto-encoder architecture, where the SPV representations can be seen as a latent encoding of the original input (or some transformation thereof) [24]. In this view, the efficient encoding of the rather complex visual environment into a relatively rudimentary phosphene representation can be considered a dimensionality reduction problem in which we aim to maximally preserve the information that is present in the latent SPV representation.

**Table 2.**
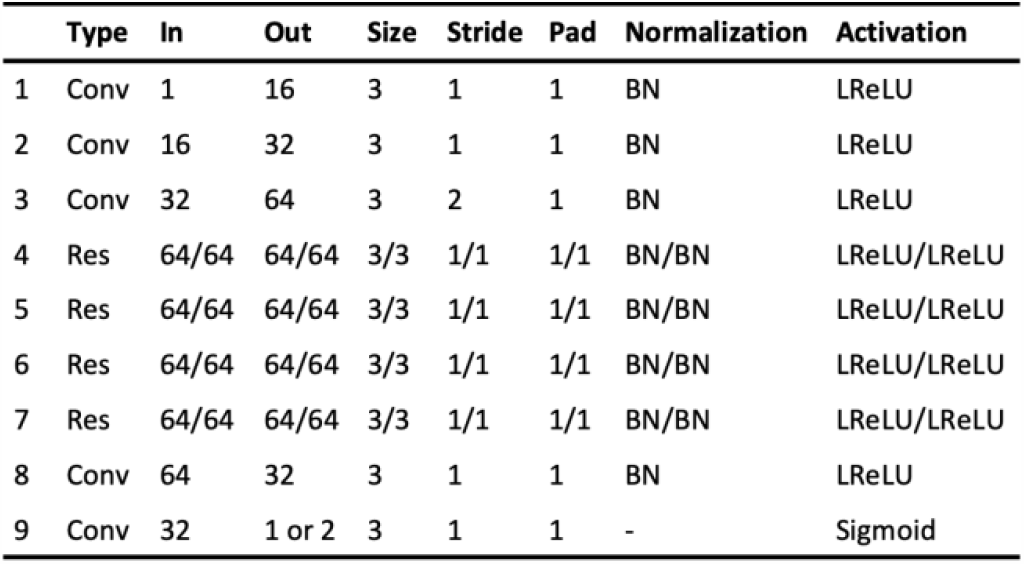
Architecture of the encoder component. Conv: convolutional layer; Res: residual block; Pool: max-pooling layer; BN: batch normalization; LReLU: leaky rectified linear unit.

## Experiments and Results

Model performance was explored via three computational experiments using different datasets. In each experiment a different combination of loss functions and constraints was tested, as explained below. To quantify reconstruction performance, we report suitable image quality assessment measures. Unless stated otherwise, we report the mean squared error (MSE), the structural similarity index (SSIM) and either the peak signal to noise ratio (PSNR) or the feature similarity index (FSIM) between the reconstruction and the input image [25, 26]. In addition, we report the average percentage of activated electrodes. Furthermore, to visualize the subjective quality of encodings and reconstructions, we display a subset of images with the corresponding SPV representations and image reconstructions.

### Training procedure

The end-to-end model was implemented in PyTorch version 1.3.1., using a NVIDIA GeForce GTX 1080 TI graphics processing unit (GPU) with CUDA driver version 10.2. The trainable parameters of our model were updated using the Adam optimizer [27]. To account for potential convergence of the model parameters towards local optima (i.e. to reduce the likelihood that a suitable parameter configuration of the network is missed due to a combination of a specific weights initialization and learning rate), a ‘random restarts’ approach was used. That is, each model was trained five times, whilst each time randomly starting with a different weight initialization. The results only show the best performing one out of these five models (i.e. the one with the lowest loss on the validation dataset), unless stated otherwise.

### Experiment 1: SPV-based reconstruction of visual stimuli

The objective of the first experiment, was to test the basic ability of the proposed end-to-end model to encode a stimulus into regular, binary phosphene representations, and decode these into accurate reconstructions of the input image. For this purpose, we trained the model on a self-generated dataset containing basic black and white images with a randomly positioned lowercase alphabetic character. Each image in the dataset is 128 × 128 pixels, and contains a randomly selected character, displayed in one of 47 fonts (38 fonts for the training dataset, 9 fonts for the validation dataset). The model was trained to minimize the (pixel-wise) mean squared error

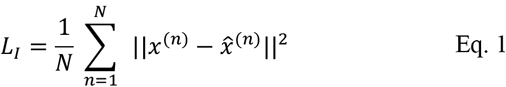

between the intensities of the input image *x*^(*n*)^ and the output reconstruction 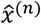 over all training examples 1 ≤ *n* ≤ *N*.

The results of experiment 1 for the best performing model out of the five random restarts are displayed in figure 3. As can be observed, the reconstruction loss is successfully minimized until convergence after 71 epochs. The performance metrics on the validation dataset indicate that the model is capable of adequately reconstructing the input image from the generated SPV representation (MSE = 0.007, SSIM = 0.693, PSNR = 21.99). Notably, the model has adopted an encoding strategy where the presence of the alphabetic character (white pixels) is encoded with the absence of phosphenes and vice versa, resulting in an average electrode activity (percentage of active electrodes) of 90.69%.

**Figure 3.**
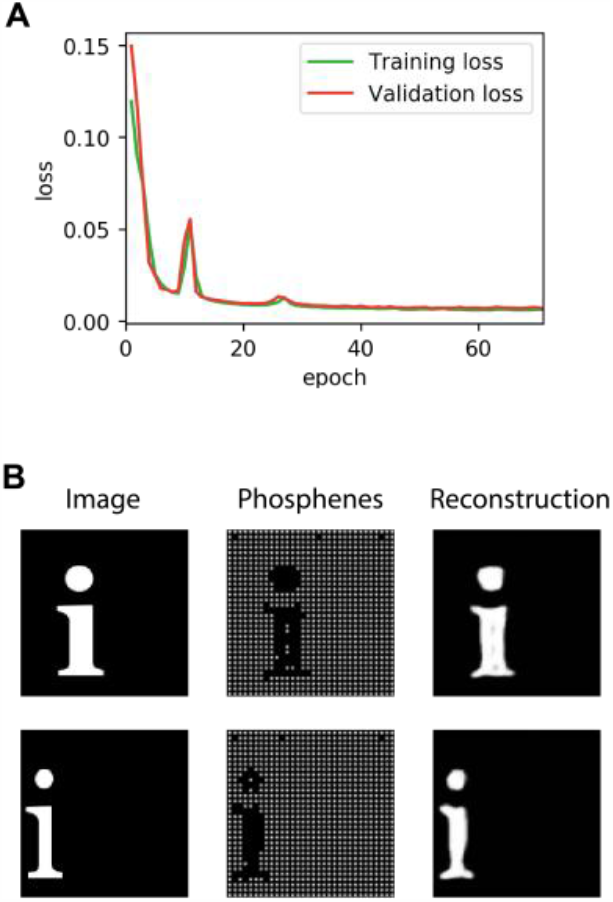
Results of experiment 1. The model was trained to minimize mean squared error loss. A. Visualization of the network input (left) the simulated prosthetic vision (middle) and the reconstruction (right). B. The training curves indicating the loss on the training dataset and validation dataset during the training procedure.

### Experiment 2: Shaping phosphene vision via constrained optimization

In the second experiment, we assess whether our model allows the inclusion of additional constraints. To exemplify this advantage of our proposed approach, we chose to evaluate the effects of adding a sparsity loss term. Considering the potentially adverse effects of electrical stimulation [5, 28], one might want to introduce such a sparsity requirement that constrains the stimulation protocol, limiting the neural degradation by enforcing minimal energy transfer to neural tissue. Let *s* = (*s*_1_, …, *s*_*M*_) denote a stimulation vector representing the stimulation pattern for *M* electrodes. We define sparsity as the L1 norm on the stimulation protocol:

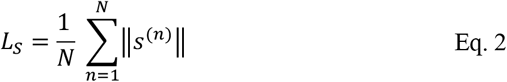

where *s*^(*n*)^ is the stimulation vector for the *n*-th training example. The objective is to minimize the total loss:

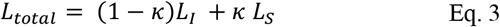

where *L*_*I*_ is the pixel-wise reconstruction loss and the parameter *κ* can be adjusted to choose the relative weight between the reconstruction loss and sparsity loss. We evaluated 13 values of *κ*, again each time using the random restarts approach, testing five different weight initializations. We performed a regression analysis on the overall percentage of active electrodes and the reconstruction performance (MSE) to evaluate the effectiveness and the decrease in performance, respectively.

The results for three of the κ-parameters are displayed in figure 4. As can be observed, adding additional sparsity loss by increasing the κ parameter, resulted in an overall lower percentage of active phosphenes. For larger values of κ, the reconstruction quality dropped. In contrast to the results of experiment 1, addition of a sparsity loss led to phosphene patterns that more naturally encode the presence of pixels by the presence of phosphenes (instead of vice versa).

**Figure 4.**
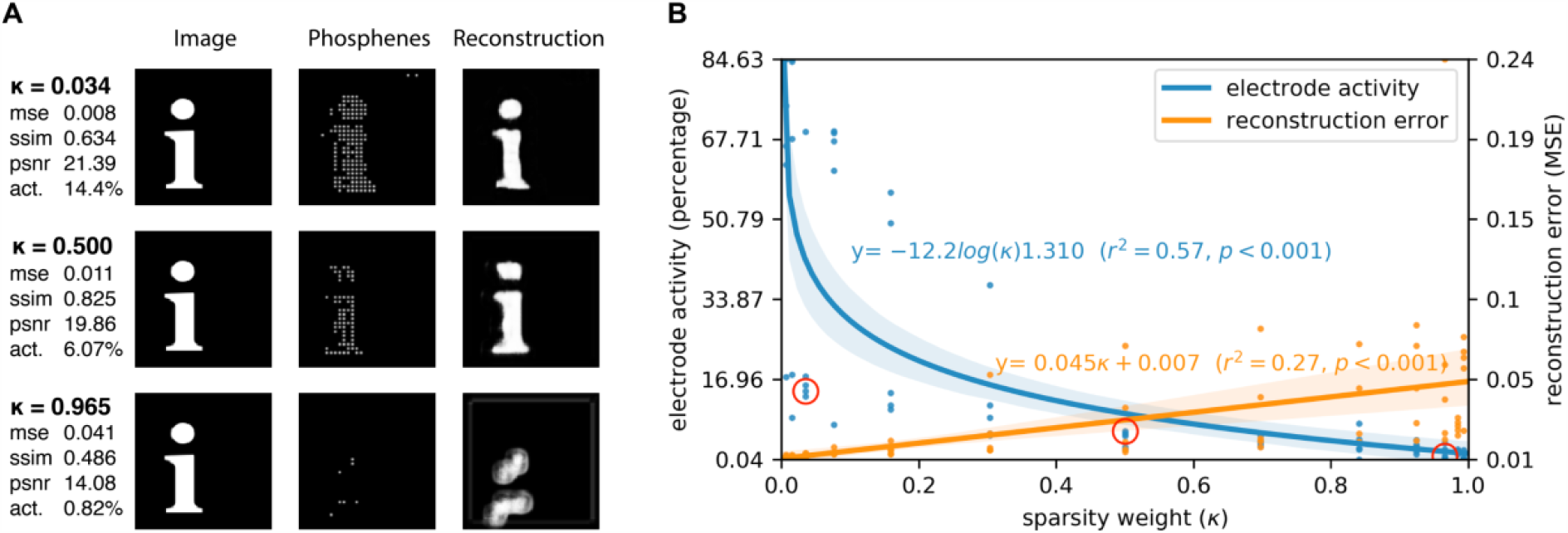
Results of experiment 2. The model was trained on a combination of mean squared error loss and sparsity loss. 13 different values for sparsity weight *k* were tested. A. Visualization of the results for three out of the 13 values for *k*. Each row displays the performance metrics for the best-performing model out of five random restarts, and one input image from, the validation dataset (left), with the corresponding simulated phosphene representation (middle) and reconstruction (right). B. Regression plot displaying the sparsity of electrode activation and the reconstruction error in relation to the sparsity weight *k*.

### Experiment 3: End-to-end phosphene vision for naturalistic tasks

In the last experiment, the model was trained on a more complex and naturalistic image dataset. To this end, we made use of the ADE20k semantic segmentation dataset [29, 30]. Compared with the synthetic character dataset which we used for the aforementioned experiments, one of the key challenges of such naturalistic stimuli is that instead of merely foreground objects on a plain dark background, the images contain abundant information. Here, not all features may be considered relevant, and therefore the task at hand (implemented by a loss function) should control which information needs to be preserved in the phosphene representations. To explore different strategies for solving such a challenge, we compared the pixel-based MSE reconstruction task that was used in the first experiment with two other types of reconstruction tasks: First, an unsupervised perceptual reconstruction task, similar to ref. [31, 32], which, in contrast to the pixel-wise MSE-based reconstruction task, aims to only preserve high-level perceptual features. Second, a supervised semantic boundary reconstruction task, to evaluate the additional value of using labeled supervision to specify which information needs to remain preserved in the reconstructions.

The perceptual reconstruction task was formulated with the aim of minimizing higher-level perceptual differences between the reconstruction and the input image. These more abstract perceptual differences are defined in feature-space, as opposed to the more explicit per-pixel differences which were used in the previous experiments. The feature loss is defined as

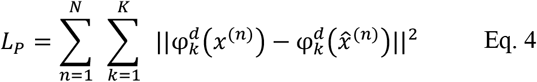

where N is the number of training examples, φ^d^(*x*^(*n*)^) and 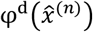 are the *d*-th layer feature maps extracted from the input image and the reconstruction image using the VGG16 model pre-trained on the ImageNet dataset [33] and *K* is the number of feature maps of that layer. For lower values of *d*, the perceptual loss reveals more explicit differences in low-level features such as intensities and edges, whereas for higher values of *d* the perceptual loss focuses on more abstract textures or conceptual differences between the input and reconstruction [34]. We chose *d* equal to 3 as an optimal depth for the feature loss, based on a comparison between different values (see figure 5).

**Figure 5.**
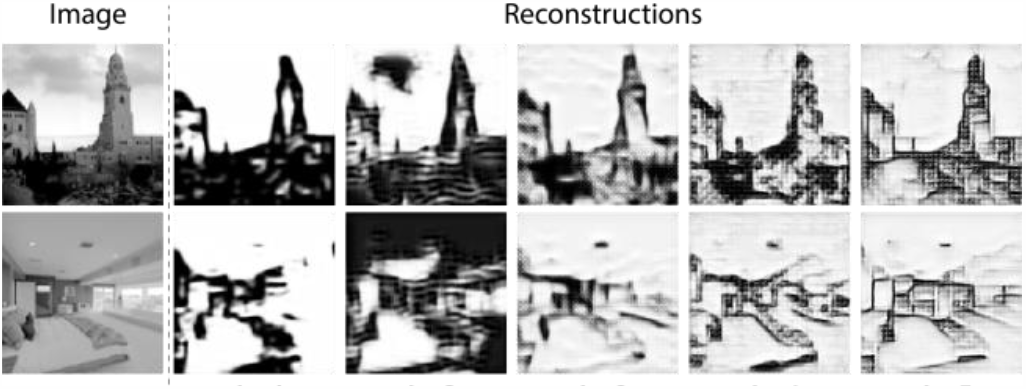
Comparison between different values of d for the perceptual reconstruction task that was used in experiment 3, where d indicates the layer depth for the VGG-based feature loss.

In the supervised semantic boundary reconstruction task, the objective was not to minimize the differences between reconstruction and input image. Instead, we aimed to provide labeled supervision that guides the model towards preserving information of semantically defined boundaries. Here, the objective was to minimize the differences between the output prediction of the decoder and processed semantic segmentation target labels of the ADE20K dataset. The reconstruction loss was formalized as the weighted binary cross entropy (BCE), defined as

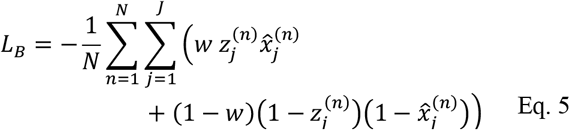

where 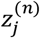 is the ground truth boundary segmentation label for pixel *j* of example *n* and *w* is a constant that is introduced for counterbalancing the unequal distribution of non-boundary compared to boundary pixels. In our experiments w is set equal to 0.925, matching the inverse ratio of boundary pixels to non-boundary pixels. Both for the perceptual loss and the BCE loss we included a sparsity loss as in experiment 2.

The results of experiment 3 are displayed in figure 6 and table 3. The performance metrics of the intensity-based reconstruction task (MSE = 0.032, SSIM = 0.574) were lower compared to those on the basic character dataset that was used in experiment 2. Both for the perceptual reconstruction tasks and the supervised semantic boundary reconstruction task, the model seems to have adopted a different phosphene encoding strategy compared to the intensity-based reconstruction task. The average MSE in the perceptual reconstruction task is higher, but the FSIM was lower compared to the intensity-based reconstruction task. In the supervised semantic boundary condition, 65.6% of the boundary pixels were classified correctly. The average percentage of active electrodes was lowest in this condition, compared with the other two conditions.

**Table 3.** Performance metrics for experiment 3. Act.: percentage of activated electrodes; MSE: mean squared error; FSIM: feature similarity index; SSIM: structural similarity index; Acc.: Accuracy; Sens.: sensitivity; Spec.: specificity.

| Reconstruction Task | Act. (%) | MSE | FSIM | SSIM | Acc. | Sens. | Spec. |
| --- | --- | --- | --- | --- | --- | --- | --- |
| Intensity-based | 59.94 | 0.032 | 0.723 | 0.574 | - | - | - |
| Perceptual | 41.52 | 0.059 | 0.767 | 0.556 | - | - | - |
| Semantic boundary | 36.59 | - | - | - | 0.656 | 0.800 | 0.644 |

**Figure 6.**
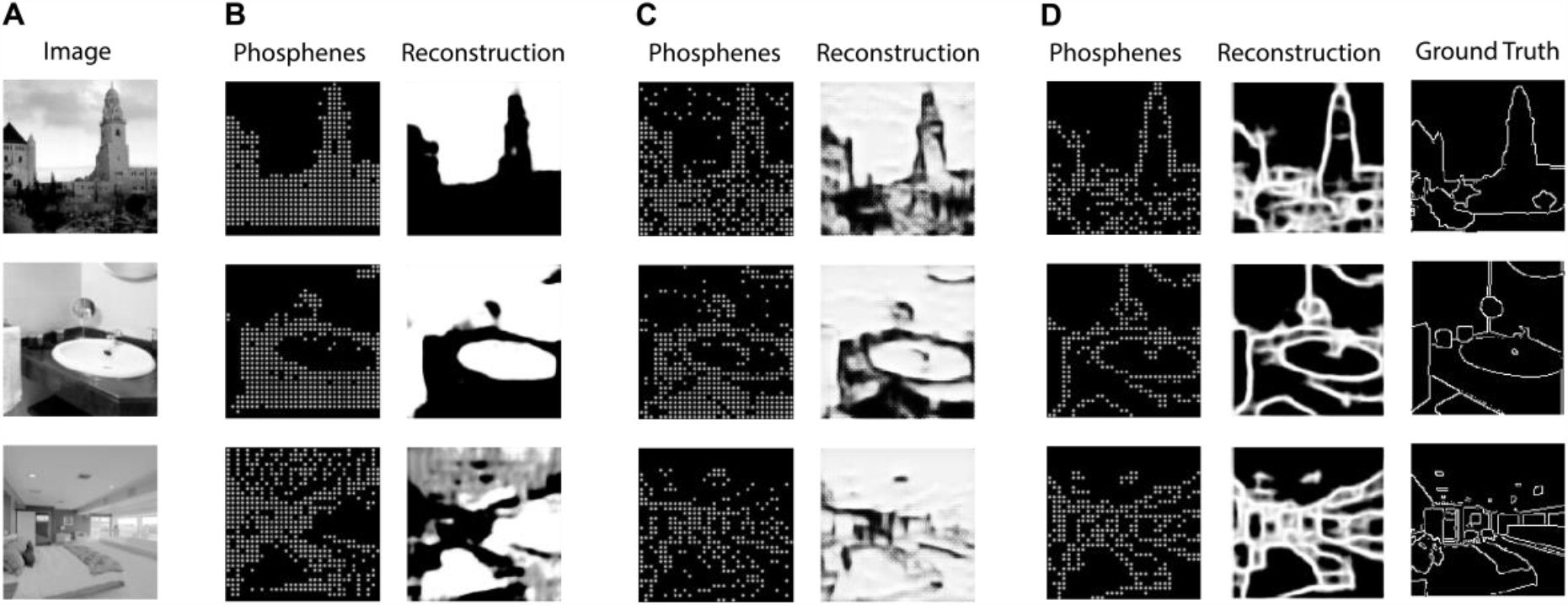
Results of experiment 3. The model was trained on naturalistic stimuli, comparing three reconstruction tasks. A) original image B) Pixel intensity-based reconstruction task with MSE loss C) Perceptual reconstruction task, using VGG feature loss (d is set equal to 3) D) Semantic boundary reconstruction task, using weighted BCE loss between the reconstruction and the ground truth label that is provided in the dataset.

## Discussion

In this paper we present and evaluate a novel DL approach for end-to-end optimization of prosthetic vision. Below we provide a general discussion of the proposed method and the results of our validation experiments, reflecting on the earlier hypothesized automated and tailored optimization abilities. Furthermore, we list some of the limitations of the current study and provide directions for future research.

### Automated optimization

Our end-to-end model is based on an autoencoder architecture, and aims to make use of their well-described ability to efficiently encode information into a low-dimensional latent representation [24]. The results from experiment 1 demonstrate that the model successfully converges to an optimal encoding strategy for a latent representation that consisted of a 32 × 32 binary simulated phosphene pattern. The model achieved adequate reconstruction performance, as indicated by the low MSE of 0.007 on the validation dataset. Note, that in the unconstrained setting, the possible phosphene encoding strategies that may be found by the model are not limited to practical, ecologically useful, solutions. This is exemplified by the large average electrode activity of 90.69%. To guide the model towards useful latent SPV representations, in experiments 2 and 3 we implement additional regularizing constraints (sparsity), and explore different optimization tasks, as discussed below.

An important requirement for automated optimization with deep learning, is that all components of the artificial neural network make use of differentiable operations. In this paper we contribute a basic implementation of a fully differentiable phosphene simulation module, including a straight-through estimator for quantized phosphene activation. Further studies could adapt or extend this implementation to test automated optimization for different phosphene simulations, for instance, varying the number of electrodes, and the positions of the phosphenes.

### Tailored optimization to sparsity constraints

Implementing additional constraints in the optimization procedure may provide a general solution to account for practical, medical or biophysical limitations of the prosthetic device. For instance, the inflammatory response of brain tissue is a major concern that limits the long-term viability of cortical electrode implants [35, 36] and these adverse effects may partly be avoided by limiting the chronic electrical stimulation itself [5, 28]. The results of experiment 2 demonstrate that implementation of a such a sparsity constraint may help to regularize the electrode activity. Notably, a larger sparsity weight *κ* results in fewer active electrodes, but also in impaired reconstruction performance. Choosing a balanced value for *κ*, depending on the needs of the patient, can be seen as a part of a tailored optimization approach of image preprocessing in prosthetic vision.

Importantly, the proposed method enables the implementation of virtually any type of additional constraint that can be incorporated in the optimization procedure. Other examples of biophysical limitations for prosthetic vision, besides sparse electrode activation, could include minimal distance for simultaneously activated electrodes, maximal spread of electrode use or minimal temporal separation. Future research focusing on such biophysical limitations could extend the proposed method to include such or other constraints in the optimization procedure.

### Task-specific optimization for naturalistic settings

Due to the relative complexity, and the presence of non-relevant information, the encoding of naturalistic scenes into phosphenes remains a challenge and it requires task-dependent processing. This challenge is explicitly addressed by the proposed end-to-end approach.

The results of experiment 3 show that for different tasks the model converges to a different optimal encoding strategy, which may indicate task-specific optimization. The higher FSIM and the lower MSE in the perceptual reconstruction task, compared to the intensity-based reconstruction task, indicate that information about the higher-level perceptual features is favoured over pixel-intensity information. Similarly, when the model was trained with BCE-loss to reconstruct the processed target labels from the ADE20k dataset, only semantic boundary information was preserved.

The phosphene encodings that were found by the model in this condition show similarities to those in ref. [37], who demonstrated that pre-processing with semantic segmentation may successfully improve object recognition performance in simulated prosthetic vision (compared to pre-processing with conventional edge detection techniques). Note, that in the proposed end-to-end architecture, supervised training was merely applied to the output reconstructions and that the labels do not directly control the phosphene representations themselves. This method combines the advantages of existing deep learning approaches using large precisely-labeled datasets with those of tailored optimization to additional constrains, such as sparsity.

The VGG-feature loss and BCE-loss that were implemented in this paper, were not chosen only because of their well-established application in optimization problems [38, 39] but also because they represent basic functions that are normally performed in the brain. The feature representations found in deep neural networks illustrate a similar processing hierarchy to that of the visual cortex [40, 41] and boundary detection is one of these processing steps, needed for segregation of objects from background [42]. Although many details about the downstream information processing of direct stimulation in V1 are yet to be discovered, we know that conscious awareness of a stimulated percept requires coordinated activity across a whole network of brain areas [43]. By acting as a digital twin, a well-chosen reconstruction task may mimic the downstream visual processing hierarchy, enabling direct optimization of visual prosthetics to the biological system. Still, fully optimizing the interaction between prosthetic stimulation and the downstream visual processing, requires a deep understanding of the biological networks involved. The proposed end-to-end approach is designed in a modular way and future research can extend the concept with virtually any reconstruction model and task.

### Limitations and future directions

Some limitations of the present study provide directions for future research. First, the subjective quality of the phosphene representations is not addressed in the current study. Future research could compare the phosphene encoding strategies found by our proposed model, to existing pre-processing approaches from the current literature, using behavioural experiments. Second, the simulated prosthetic vision that was used in the current study is a simplified model of the reality. Future research could extend the current model to simulate more realistic features of phosphene perception, such as different shapes, locations and sizes of the phosphenes. Third, in this paper the model is trained on static images. Future approaches could extend our end-to-end model to process dynamical stimuli, resembling an even more naturalistic setting. Finally, the optimization tasks that were used in the current paper, remain basic. Future work could extend the current approach with other or more complex tasks. For instance, with reinforcement learning strategies, the model could be extended to perform tasks that more closely related to the everyday actions that need to be performed by the end-user, such as object manipulation [44] or object avoidance [45].

## Conclusion

In this paper we present a novel DL-based approach for automated and tailored end-to-end optimization of prosthetic vision. Our validation experiments show that such an approach may help to automatically find a task-specific stimulation protocol, considering an additional sparsity requirement. The presented approach is highly modular and could be extended to dynamically optimize prosthetic vision for everyday tasks and requirements of the end-user.

## Notes

### Competing Interest Statement

The authors have declared no competing interest.

